# gwasurvivr: an R package for genome wide survival analysis

**DOI:** 10.1101/326033

**Authors:** Abbas A Rizvi, Ezgi Karaesmen, Martin Morgan, Leah Preus, Junke Wang, Michael Sovic, Theresa Hahn, Lara E Sucheston-Campbell

## Abstract

**Summary:** To address the limited software options for performing survival analyses with millions of SNPs, we developed gwasurvivr, an R/Bioconductor package with a simple interface for conducting genome wide survival analyses using VCF (outputted from Michigan or Sanger imputation servers), IMPUTE2 or PLINK files. To decrease the number of iterations needed for convergence when optimizing the parameter estimates in the Cox model we modified the R package survival; covariates in the model are first fit without the SNP, and those parameter estimates are used as initial points. We benchmarked gwasurvivr with other software capable of conducting genome wide survival analysis (genipe, SurvivalGWAS_SV, and GWASTools). gwasurvivr is significantly faster and shows better scalability as sample size, number of SNPs and number of covariates increases.

**Availability and implementation:** gwasurvivr, including source code, documentation, and vignette are available at: http://bioconductor.org/packages/gwasurvivr

**Contact:** Abbas Rizvi; Lara E Sucheston-Campbell

Supplementary information: Supplementary data are available at https://github.com/suchestoncampbelllab/gwasurvivr_manuscript

## 1 Introduction

Genome-wide association studies (GWAS) are population-level experiments that investigate genetic variation in individuals to observe single nucleotide polymorphism (SNP) associations with a phenotype. Genetic variants tested for association are genotyped on an array and imputed from a reference panel of sequenced genomes, e.g. 1000 Genomes Project or Haplotype Reference Consortium (HRC) (Das, et al., 2016; Genomes Project, et al., 2015). Imputation increases genome coverage from hundreds of thousands or a few million to upwards of 30 million SNPs, improves power to detect genetic associations, and/or homogenizes variant sets for meta-analyses (Das, et al., 2016). Imputed SNPs can be tested for association with binary outcomes (case/control) and quantitative outcomes (e.g., height) using a range of available software packages, including SNPTEST (Marchini, et al., 2007) or PLINK (Purcell, et al., 2007). However, existing software options for performing survival analyses, genipe (Lemieux Perreault, et al., 2016), SurvivalGWAS_SV (Syed, et al., 2017), and GWASTools (Gogarten, et al., 2012) either require user interaction with raw output, were not initially designed for survival and/or have long run times. For these reasons, we developed an R/Bioconductor package, gwasurvivr, for genome wide survival analyses of imputed data in multiple file formats with flexible analysis and output options.

## 2 Implementation

### 2.1 Data structure

Gwasurvivr can analyze data in IMPUTE2 format (Howie, et al., 2009), in VCF files derived from Michigan (Das, et al., 2016) or Sanger imputation servers (McCarthy, et al., 2016), and directly genotyped PLINK format (Purcell, et al., 2007). Data from each are prepared in gwasurvivr by leveraging existing Bioconductor packages GWASTools (Gogarten, et al., 2012) or VariantAnnotation (Obenchain, et al., 2014) depending on the imputation file format.

#### IMPUTE2 FORMAT

IMPUTE2 (Howie, et al., 2009) format is a standard genotype (.gen) file which store genotype probabilities (GP). We utilized GWASTools in R to compress files into genomic data structure (GDS) format (Gogarten, et al., 2012). This allows for efficient, iterative access to subsets of the data, while simultaneously converting GP into dosages (DS) for use in survival analyses.

#### VCF

VCF files generated from these Michigan or Sanger servers include a DS field and server-specific meta-fields (INFO score [Sanger] or r^2^ [Michigan], as well as reference panel allele frequencies) that are iteratively read in by VariantAnnotation (Obenchain, et al., 2014).

#### PLINK FORMAT

Plink bed files contain genotype information encoded in binary format. Fam and bim files include the information of phenotype and marker location, respectively (Purcell, et al., 2007).

### 2.2 Survival analysis

gwasurvivr implements a Cox proportional hazards regression model (Cox, 1992) to test each SNP with an outcome with options for including covariates and/or SNP-covariate interactions. To decrease the number of iterations needed for convergence when optimizing the parameter estimates in the Cox model we modified the R package survival (Therneau and Grambsch, 2000). Covariates in the model are first fit without the SNP, and those parameter estimates are used as initial points for analyses with each SNP. If no additional covariates are added to the model, the parameter estimation optimization begins with null initial value. (Supplementary Figure 1).

Survival analyses are run using genetic data in either VCF or IMPUTE2 (Howie, et al., 2009) formats and a phenotype file, which contains survival time, survival status and additional covariates, both files are indexed by sample ID. In addition to genomic data, the VCF files contain both sample IDs and imputation quality metrics (INFO score or r^2^), while IMPUTE2 (Howie, et al., 2009) come in separate files (.gen,.sample, and .info).

Gwasurvivr functions for IMPUTE2 (impute2CoxSurv or gdsCoxSurv) and VCF (michiganCoxSurv or sangerCoxSurv) include arguments for the survival model (event of interest, time to event, and covariates) and arguments for quality control that filter on minor allele frequency (MAF) or imputation quality (michiganCoxSurv and sangerCoxSurv only). INFO score filtering using impute2CoxSurv can be performed by accessing the .info file from IMPUTE2 results and subsequently providing the list of SNPs to ‘exclude.snps’ argument to gwasurvivr. Users can also provide a list of sample IDs for gwasurvivr to internally subset the data. gwasurvivr outputs two files: (1).snps_removed file, listing all SNPs that failed QC parameters and (2).coxph file with the results from the analyses, including parameter estimates, p-values, MAF, the number of events and total sample N for each SNP. gwasurvivr also allows the number of cores used during computation on Windows and Linux to be specified. Users can keep compressed GDS files after the initial run by setting keepGDS argument to TRUE when analyzing IMPUTE2 data (Howie, et al., 2009). On successive runs, gdsCoxSurv can then be used instead of impute2CoxSurv to avoid compressing the data on each GWAS run.

## 3 Simulations and benchmarking

Computational runtimes for gwasurvivr were benchmarked against existing software comparing varying sample sizes and SNP numbers, with 4, 8 or 12 covariates and for a single chromosome with 15,000-25,000 individuals. In addition, we evaluated time for gwasurvivr for a GWAS (~6 million SNPS) for 3000, 6000 and 9000 samples. All benchmarking experiments were performed using IMPUTE2 format (comparison packages do not take VCF from either imputation servers).

Description of simulated genotype and phenotype data are in the Supplementary Data.

## 4 Results

gwasurvivr was faster than genipe (Lemieux Perreault, et al., 2016), SurvivalGWAS_SV (Syed, et al., 2017), and GWASTools (Gogarten, et al., 2012) for 100,000 SNPs at N=100, and 5000, with the exception of SurvivalGWAS_SV at n=1000 (**Figure 1A**). Similarly, increasing the number of covariates for gwasurvivr has minimal effects on runtime versus other software (**Figure 1B**). Gwasurvivr computes for large sample sizes, however, compression time increases with increasing sample size, and likely will be limited by available RAM on a machine or cluster (**Figure 1C**). The keepGDS argument helps address this and results in reduced run times (**Figures 1C** and **1D**), i.e. <3 hours for a GWAS of N=9,000. A ~6 million SNP GWAS can be run in <10 hours for 9000 samples when using separately scheduled jobs on a supercomputer (**Figure 1D**). However, gwasurvivr overcomes memory limitations often attributed to R by processing subsets of the entire data, and thus it is possible to conduct genome-wide survival analyses on a typical laptop computer.

**Figure 1.**
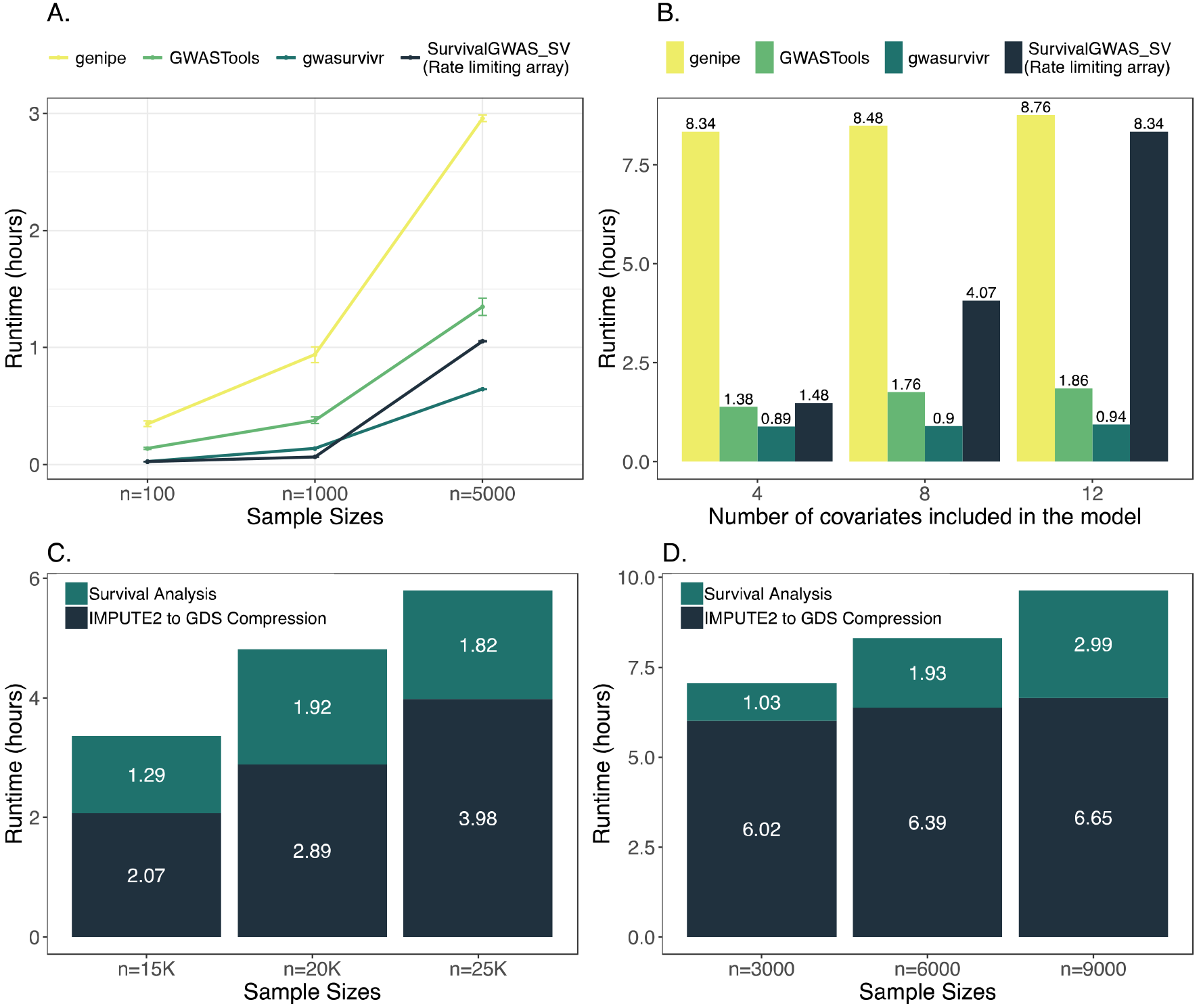
Runtime for survival analyses. All analyses were run with identical CPU constraints of 1 node and 8 cores. All SurvivalGWAS_SV runtimes are shown for the rate limiting array for 100 batched jobs with 1,000 SNPs in an array index. The run time for the rate-limiting array is defined as the array index that had the longest runtime. This gives the shortest possible runtime for SurvivalGWAS_SV where all submitted jobs start at the same time, without pending in the cluster queue. **A.** The x-axis shows the three sample sizes (n=100, 1000, 5000) with 100,000 SNPs. The y-axis is the total runtime in hours. The mean runtime and 95% confidence intervals (CI) are show for genipe (yellow), GWASTools (light green), SurvivalGWAS_SV (dark blue) and gwasurvivr (dark green). Confidence intervals were calculated for three simulations for each n and m combination. **B.** Genipe (yellow), GWASTools (light green), SurvivalGWAS_SV (dark blue), and gwasurvivr (dark green) run with 4, 8, and 12 covariates (n=5000, m=100,000). **C.** Gwasurvivr was run on IMPUTE2 data simulated from chromosome 22 (m~117,000 SNPs) for n=15,000, n=20,000 and n=25,000. The dark blue shaded area is elapsed time for compressing IMPUTE2 to GDS format and the dark green shaded areas are the computational time to run the survival analysis alone. **D.** Full GWAS were run using gwasurvivr on sample sizes of n=3000, n=6000, and n=9000. Shown here is the amount of time for the largest chromosome (chromosome 2) to complete. This corresponds to the full time for a GWAS when using a job scheduler on a cluster. The dark blue shaded area is elapsed time for compressing IMPUTE2 to GDS format andthe dark green shaded areas are the computational time to run the survival analysis alone. Supplementary Figure 5 contains the runtime for all chromosomes.

gwasurvivr is a fast, efficient, and flexible program well suited for multi-core processors and easily run in a computing cluster environment.

## Funding

This work was supported by the National Institutes of Health/National Heart, Lung, and Blood Institute R01HL102278 [to LSC and TH], National Institutes of Health/National Cancer Institute [R03CA188733], The Ohio State University and the Translational Data Analytics Initiative, and the Pelotonia Graduate Student Fellowship [to EK]

## ACKNOWLEDGEMENTS

The authors would like to thank Amy Webb and Guy Brock at The Ohio State University Department of Biomedical Informatics for listening to presentations on and offering suggestions about gwasurvivr. This work was performed in part at the University at Buffalo’s Center for Computational Research.

## REFERENCES

Cox, D.R. Regression Models and Life-Tables. Springer; 1992.

Das, S., et al. Next-generation genotype imputation service and methods. Nat Genet 2016;48(10):1284–1287.

Genomes Project, C., et al. A global reference for human genetic variation. Nature 2015;526(7571):68–74.

Gogarten, S.M., et al. GWASTools: an R/Bioconductor package for quality control and analysis of genome-wide association studies. Bioinformatics 2012;28(24):3329–3331.

Howie, B.N., Donnelly, P. and Marchini, J. A flexible and accurate genotype imputation method for the next generation of genome-wide association studies. PLoS Genet 2009;5(6):e1000529.

Lemieux Perreault, L.P., et al. genipe: an automated genome-wide imputation pipeline with automatic reporting and statistical tools. Bioinformatics 2016;32(23):3661–3663.

McCarthy, S et al. (2016) A reference panel of 64,976 haplotypes for genotype imputation, Nature Genetics. 48(10):1279–83 [27548312]

Marchini, J., et al. A new multipoint method for genome-wide association studies by imputation of genotypes. Nat Genet 2007;39(7):906–913.

Obenchain, V., et al. VariantAnnotation: a Bioconductor package for exploration and annotation of genetic variants. Bioinformatics 2014;30(14):2076–2078.

Purcell, S., et al. PLINK: a tool set for whole-genome association and population-based linkage analyses. Am J Hum Genet 2007;81(3):559–575.

Su, Z., Marchini, J. and Donnelly, P. HAPGEN2: simulation of multiple disease SNPs. Bioinformatics 2011;27(16):2304–2305.

Syed, H., Jorgensen, A.L. and Morris, A.P. SurvivalGWAS_SV: software for the analysis of genome-wide association studies of imputed genotypes with “time-to-event” outcomes. BMC Bioinformatics 2017;18(1):265.

Therneau, T.M. and Grambsch, P.M. Modeling Survival Data: Extending the Cox Model. Springer; 2000.

